# Cystathionine γ Lyase Deletion Enhances Corpus Cavernosum Contraction via Thromboxane A_2_ and Neurogenic Pathways Without Affecting Endothelial Function

**DOI:** 10.1101/2025.05.17.654668

**Authors:** Tooyib A. Azeez, Clifford J. Pierre, Colin M. Ihrig, Stephen P. Chelko, Judy M. Muller-Delp, Justin D. La Favor

## Abstract

Cystathionine γ-lyase (CSE) produces hydrogen sulfide (H₂S), a vasodilator critical for vascular function. While its systemic effects are well-documented, its role in erectile physiology remains unclear. This study investigated the impact of CSE deletion on vascular and erectile tissue reactivity. We hypothesized that CSE knockout (CSE-KO) mice would exhibit endothelial dysfunction. A total of 22 CSE-KO and 22 age-matched wild-type (WT) controls were studied at one year of age. The internal iliac artery (IIA), internal pudendal artery (IPA), and corpus cavernosum (CC) were harvested for *ex vivo* functional assessments using tissue, wire, and pressure myography. Vasoconstriction was evaluated using phenylephrine, endothelin-1, U-46619, and electrical field stimulation (EFS). Endothelium-dependent relaxation was assessed using acetylcholine (ACh) and flow-mediated dilation, while endothelium-independent relaxation was evaluated using sodium nitroprusside (SNP). Sodium sulfide (Na₂S) was used to assess H₂S-mediated dilation. Non-adrenergic, non-cholinergic (NANC) transmission was evaluated using EFS. No significant differences were observed in ACh-, SNP-, or flow-mediated relaxation, although CSE-KO mice demonstrated impaired NANC-nerve mediated relaxation in the CC. Moreover, CSE-KO mice exhibited significantly enhanced CC contraction in response to U-46619 and EFS, suggesting increased vascular resistance in the end organ CC rather than the pre-penile arteries. Histological analysis revealed no significant structural or fibrotic remodeling in any tissue, although there was a trend toward increased collagen deposition in the IIA and IPA. These findings indicate that chronic CSE deficiency does not impair endothelial function but alters neurogenic control and increases vasoconstrictive sensitivity specifically in the CC, potentially predisposing to erectile dysfunction.

**NEW & NOTEWORTHY:** This study highlights the critical role of hydrogen sulfide (H₂S) in erectile physiology by demonstrating that CSE deletion does not impair endothelial function but significantly enhances neurogenic and thromboxane A2 receptor-induced vasoconstriction specifically in the corpus cavernosum (CC). These findings suggest that endogenous H₂S modulates neurovascular control of erection. Its deficiency predisposes the erectile system to heightened vascular resistance predominantly in the end organ, providing novel insights into the vascular mechanisms underlying erectile dysfunction.

## INTRODUCTION

Erectile dysfunction (ED) is a prevalent disorder affecting approximately 30-50% of men over the age of 40, with a growing body of evidence linking organic ED to cardiovascular diseases (CVD) (1, 2). ED is now recognized as an early marker of systemic vascular dysfunction, as it frequently precedes the onset of major cardiovascular events such as myocardial infarction and stroke (3, 4). The underlying pathophysiology of ED is largely attributed to endothelial dysfunction, a condition characterized by reduced nitric oxide (NO) bioavailability, increased oxidative stress, and vascular inflammation (5, 6).

Since normal erectile function is heavily reliant on vascular integrity, impairment in endothelial relaxation and smooth muscle compliance can significantly compromise penile blood flow and the erectile response (7). The erectile process is initiated through autonomic nerve activation, triggering arterial dilation in the internal iliac artery (IIA) and internal pudendal artery (IPA), leading to a rapid influx of blood into the corpus cavernosum (CC) (8). As blood accumulates, cavernous smooth muscle relaxation facilitates further engorgement, while venous outflow is restricted to sustain the erection (1). Given the regional vascular differences in erectile physiology, it is critical to investigate how vascular pathologies differentially affect pre-penile arteries (IIA and IPA) and the erectile tissue of the CC in models of vascular dysfunction.

Recent evidence has highlighted hydrogen sulfide (H₂S) as a key regulator of endothelial function, with its deficiency implicated in various cardiovascular pathologies (9–11). H₂S, an endogenous gasotransmitter produced primarily by cystathionine γ-lyase (CSE), cystathionine β-synthase (CBS), and 3-mercaptopyruvate sulfurtransferase (3-MST), functions as a potent vasodilator to modulate vascular tone, inhibit oxidative stress, and suppresses inflammatory signaling (12). Notably, studies have demonstrated that H₂S production is markedly reduced in conditions such as hypertension, atherosclerosis, and metabolic disorders, contributing to vascular stiffening, impaired vasodilation, and increased fibrosis (13, 14). In CSE knockout (CSE KO) mice, severe endothelial dysfunction and vascular remodeling have been observed across multiple vascular beds, including the aorta and mesenteric resistance arteries, suggesting that H₂S deficiency significantly exacerbates vascular dysfunction (15–18). Furthermore, emerging data indicate that H₂S plays a critical role in regulating extracellular matrix remodeling, particularly by influencing collagen deposition and smooth muscle plasticity (19, 20). These factors are essential for maintaining vascular compliance and erectile function, while fibrosis of the CC is a causative factor in ED pathogenesis in multiple pathological states (21). While the role of H₂S in systemic endothelial function has been well-studied, its impact on regional penile vasculature, particularly the IPA and CC remains unexplored, warranting further investigation into how CSE deficiency affects erectile hemodynamics.

Recent research has demonstrated suppressed CSE levels in the CC of men with severe ED, as well as rodent models of ED resulting from type 1 diabetes and a chronic Western diet (22–24). Fibrosis of the CC is a factor that drives increasing severity of ED under several pathological conditions (21, 25). While several pathways influence fibrotic progression, chronic inflammation, and oxidative stress are often linked as contributing factors (26). H2S has been identified to have anti-fibrotic effects in the cardiovascular system, many of which are attributed to the antioxidant and anti-inflammatory effects of H2S (27). Our recent work has revealed that CSE KO mice have impaired erectile function that coincides with elevated penile reactive oxygen species (ROS) (24). However, it is unclear if the functional impairment is the result of one or a combination of factors including endothelial dysfunction, smooth muscle dysfunction, neurogenic impairment, enhanced vascular resistance/vasoconstriction, or if these functional impairments are localized to the pre-penile arteries, the CC, or both. Moreover, it is unclear if the chronic excess of penile ROS resulting from CSE deletion promotes fibrosis or other remodeling in these structures that contributes to functional decline. This study aimed to examine the vascular effects of CSE deletion on erectile physiology, with particular emphasis on the IIA, IPA, and CC. We hypothesized that CSE KO mice would exhibit endothelial dysfunction in the IPA, leading to impaired vasodilation and increased vascular resistance to blood flow into the CC. Additionally, given the established role of H₂S in vascular homeostasis, we hypothesized that CSE deletion would promote fibrosis and adverse vascular remodeling, further exacerbating penile vascular dysfunction.

## MATERIALS AND METHODS

### Animal Models and Ethical Approval

All animal experiments were conducted following the Guiding Principles for the Care and Use of Vertebrate Animals in Research and Training and were approved by the Institutional Animal Care and Use Committee (IACUC) of Florida State University (Protocol Number: 202100052). Cystathionine γ-lyase knockout (CSE-KO) mice and wild-type (WT) littermate control mice were obtained in house using a heterozygous x heterozygous breeding scheme. Derivation of this strain of mice on the C57/Bl6 background has been described previously (18). A total of 44 male mice, including 22 CSE KO mice and 22 age-matched WT controls were used in this study. Mice were housed under standard conditions, provided normal chow and water *ad libitum*, and sacrificed at 12 months of age for ex vivo vascular and erectile tissue assessments. All experiments and analyses were performed by an investigator blinded to the genotype of each mouse.

### Tissue Harvesting and Preparation

Mice were anesthetized with an intraperitoneal injection of ketamine (90 mg/kg) and xylazine (10 mg/kg) and euthanized via exsanguination and double pneumothorax. All animal dissections, tissue harvest, and vascular tissue preparations were performed under dissection microscope (Leica Microsystems, Wetzlar, Germany). Suprapubic horizontal incisions were made to expose the penis, followed by a vertical incision through the abdominal wall to expose the pubic symphysis. The pubic symphysis was then bisected, and the pubic bone was cut to expose the pudendal neurovascular bundle. Muscles and ligaments were then further cleared, as described previously, to trace the dorsal penile artery at the point of entry into the penis to the internal pudendal artery (IPA), to the internal iliac artery (IIA), and to the common iliac artery (28). The distal portion of the IPA, which is distal to the gluteal artery bifurcation, was excised for study (29). A 3 mm segment of the IIA immediately proximal to the bifurcation of the proximal IPA was excised for study. After removal, artery segments were immediately placed in oxygenated ice-cold Kreb’s buffer (NaCl 130 mM, KCl 4.7 mM, KH2PO₄ 1.18 mM, MgSO₄ 1.18 mM, NaHCO₃ 14.9 mM, D-glucose 5.6 mM, CaCl₂ 1.56 mM, and EDTA 0.03 mM dissolved in distilled water) and cleaned of adherent fat and connective tissue using fine micro-spring scissors. Arteries from one side of the animal were used for vascular reactivity assessment, while arteries from the contralateral side were preserved for histology. The penis was removed from the animal at the base and the penile shaft separated from the glans penis. The dorsal vein, corpus spongiosum, and connective tissues were carefully removed from the penile shaft using fine micro-spring scissors, and the CC was bisected longitudinally along the septum for physiological reactivity assessments.

### Functional Vascular and Erectile Tissue Reactivity Assessments

To evaluate vascular function, wire and muscle strip myography were utilized to assess vasoconstriction and relaxation responses in the IIA, IPA, and CC. IIA and IPA segments (1-1.5 mm length) were mounted in a DMT 620M wire myograph system (Danish Myotechnology, Aarhus, Denmark) using 25 µm tungsten wire, while CC strips were mounted into a DMT 820MS muscle strip myograph system. Tissues were bathed in Kreb’s buffer maintained at 37 °C and continuously aerated with a 95% O2 and 5% CO2 mixture. Following 1 h of equilibration, CC tissues were stretched to a resting tension of 4 mN while artery segments were incrementally stretched and set to a resting tension corresponding to a 0.9 normalization factor (28). Following another 1 h of equilibration, tissue and vessel viability and contractile function were tested with high potassium (120 mM) Krebs buffer, with KCl substituted for NaCl. Following successive washes with Kreb’s buffer to achieve a stable resting tension, cumulative dose-response applications of vasoactive agents were initiated, as described (30). Vasoconstriction was measured in response to phenylephrine (PE, 10^-9^ to 10^-5^ M), endothelin-1 (ET-1, 10^-10^ to 10^-7^ M), and the thromboxane A2 (TXA2) receptor agonist U-46619 (10⁻⁹ to 10^-5.5^ M), with contractile responses recorded using LabChart v8 software (ADInstruments, Sydney, Australia). Endothelium-dependent relaxation was assessed using acetylcholine (ACh, 10^-9^ to 10^-5^ M), while sodium nitroprusside (SNP, 10^-9^ to 10^-5.5^ M) was used to assess endothelium-independent relaxation. Relaxation responses were performed in arteries and CC pre-constricted with 1 µM and 10 µM PE, respectively. To evaluate the effects of CSE deficiency on vascular sensitivity to hydrogen sulfide (H₂S), the rapid releasing H2S donor sodium sulfide (Na₂S, 10^-7^ to 10^-3.5^ M) was applied to pre-constricted tissues.

Neurogenic contractile responses were assessed using electrical field stimulation (EFS), where platinum electrodes delivered 20V pulses with a 2 ms pulse width for 10 s stimulations at each frequency tested, with 2 min rest between stimulations (31). Frequencies were tested over a range from 1-32 Hz. Non-adrenergic non-cholinergic (NANC) relaxation was evaluated following pretreatment with atropine (1 µM) and guanethidine (30 µM) to block cholinergic and adrenergic pathways, respectively. EFS was repeated over a frequency range of 0.5-16 Hz. EFS was applied by a CS8 stimulator (DMT) controlled by DMT MyoPULSE software. Relaxation responses were normalized as a percentage restoration to the resting tension from the PE pre-constricted value. Contractile responses were expressed as a percentage of maximal KCl-induced constriction. All vasoactive agents were dissolved in distilled water except for atropine, which was dissolved in 100% ethanol. Where atropine was added, the ethanol concentration in the bath was 0.1% (1:1000 dilution). All myograph salts and vasoactive agents were purchased from Sigma Aldrich (St. Louis, MO, USA) except for acetylcholine and U-46619, which were purchased from Enzo Biochem (New York, NY, USA).

### Pressure Myography

Pressure myography was used to assess flow-mediated dilation (FMD) responses of the IPA. IPAs were carefully cleaned as described above and placed in a physiological saline solution (PSS) composed of the following (in mmol/L): 145 NaCl, 4.7 KCl, 2.0 CaCl₂, 1.17 MgSO₄, 1.2 NaH₂PO₄, 5.0 glucose, 2.0 pyruvate, 0.02 EDTA, and 3.0 MOPS buffer, supplemented with 1 g/100 mL bovine serum albumin. The solution was adjusted to a pH of 7.4. IPA segments (1.0–1.5 mm in length) were mounted in a Lucite chamber filled with air-equilibrated PSS. Both ends of each artery were cannulated onto micropipettes with matching resistance and secured using nylon sutures. The cannulated vessels were then positioned on the stage of an inverted microscope outfitted with a video camera, a video caliper (Microcirculation Research Institute), and a data acquisition system (LabChart) to continuously monitor intraluminal diameter.

Pressurization of the vessels was achieved using two independent hydrostatic reservoirs set at 90 cmH₂O. To confirm vessel integrity, pressure was applied, and valves were subsequently closed to ensure that intraluminal diameter remained stable. Any vessels exhibiting leaks were excluded from further testing. Intact vessels were gradually warmed to 37°C and allowed to equilibrate for 30 to 60 minutes, during which spontaneous tone was allowed to develop. Once a stable basal tone was achieved, the IPAs were pre-constricted with a titration of PE until a constriction of 15-20% of the resting diameter was achieved. Flow was induced by raising and lowering paired fluid reservoirs in opposite directions, creating a pressure difference across the vessel without altering average intraluminal pressure. Changes in vessel diameter were recorded in response to pressure differentials of 2, 4, 10, 20, 40, and 60 cmH₂O (32).

### Histological Analysis of Vascular and Erectile Tissues

To investigate structural changes in vascular and erectile tissues, Masson’s Trichrome staining was performed on IIA, IPA, and CC samples to visualize collagen and smooth muscle distribution. Tissue sections were fixed in OCT compound (Sakura Finetek, Tokyo, Japan), cut into 7 µm slices, and mounted on glass slides. Sections were then sequentially exposed to Bouin’s solution, Weigert’s iron hematoxylin, Biersch Scarlet-Acid Fuchsin, phosphotungstic/phosphomolybdic acid, Aniline blue (Sigma Aldrich #HT15-1KT), and 1% acetic acid with rinses with distilled water between each exposure, as previously described (33). Stained sections were imaged using a Leica Microsystems DMi8 inverted microscope. Quantitative morphometric analyses were performed using NIH ImageJ software. Lumen area was assessed by mapping the inner wall of the artery and analyzing the area within the mapped wall. Media area was assessed by mapping the external perimeter of the arterial media and analyzing the area within the mapped wall and subtracting the lumen area. The collagen-to-smooth muscle ratio was quantified by calculating the ratio of blue-stained areas (collagen) to red-stained areas (smooth muscle) in histological sections. Histological analyses were performed in duplicate or triplicate for each tissue type from each animal and averaged values are reported.

### Statistical Analysis

All statistical analyses were performed using GraphPad Prism 9 (La Jolla, CA, USA). For ex vivo vascular reactivity assessments, two-way repeated measures ANOVA was used to compare differences between groups. Sidak’s multiple comparisons post-hoc test was used to determine group differences at individual doses where significant group differences in the ANOVA model were observed. Group differences in histological parameters were determined by unpaired t-tests. In cases where an F test determined significant differences in variances between the two groups, a non-parametric Mann-Whitney U test was used. A p-value < 0.05 was considered statistically significant.

## RESULTS

### Endothelium-Dependent and Independent Relaxation is Unaffected by CSE Deletion

To determine whether CSE deletion alters nitric oxide (NO)-mediated vasodilation, endothelium-dependent and endothelium-independent relaxation were assessed using acetylcholine (ACh) and sodium nitroprusside (SNP), respectively. Figure 1 presents cumulative dose-response curves in the internal iliac artery (IIA), internal pudendal artery (IPA), and corpus cavernosum (CC). No significant differences in ACh-induced relaxation were observed between WT and CSE-KO mice across all vascular beds (p > 0.05), suggesting that endothelial NO production remains intact. Similarly, SNP-induced relaxation did not differ significantly between groups (p > 0.05), indicating that CSE deletion does not impair smooth muscle responsiveness to NO. To further interrogate the impact of CSE deficiency on endothelial function, we assessed the dilatory response to the physiological stimulant of increasing flow in the IPAs. Flow-mediated dilation (FMD) responses are illustrated in Figure 2. The mean FMD response of the CSE KO IPA was higher across the range of flow velocity, although this trend did not reach statistical significance (p = 0.216).

**Figure 1.**
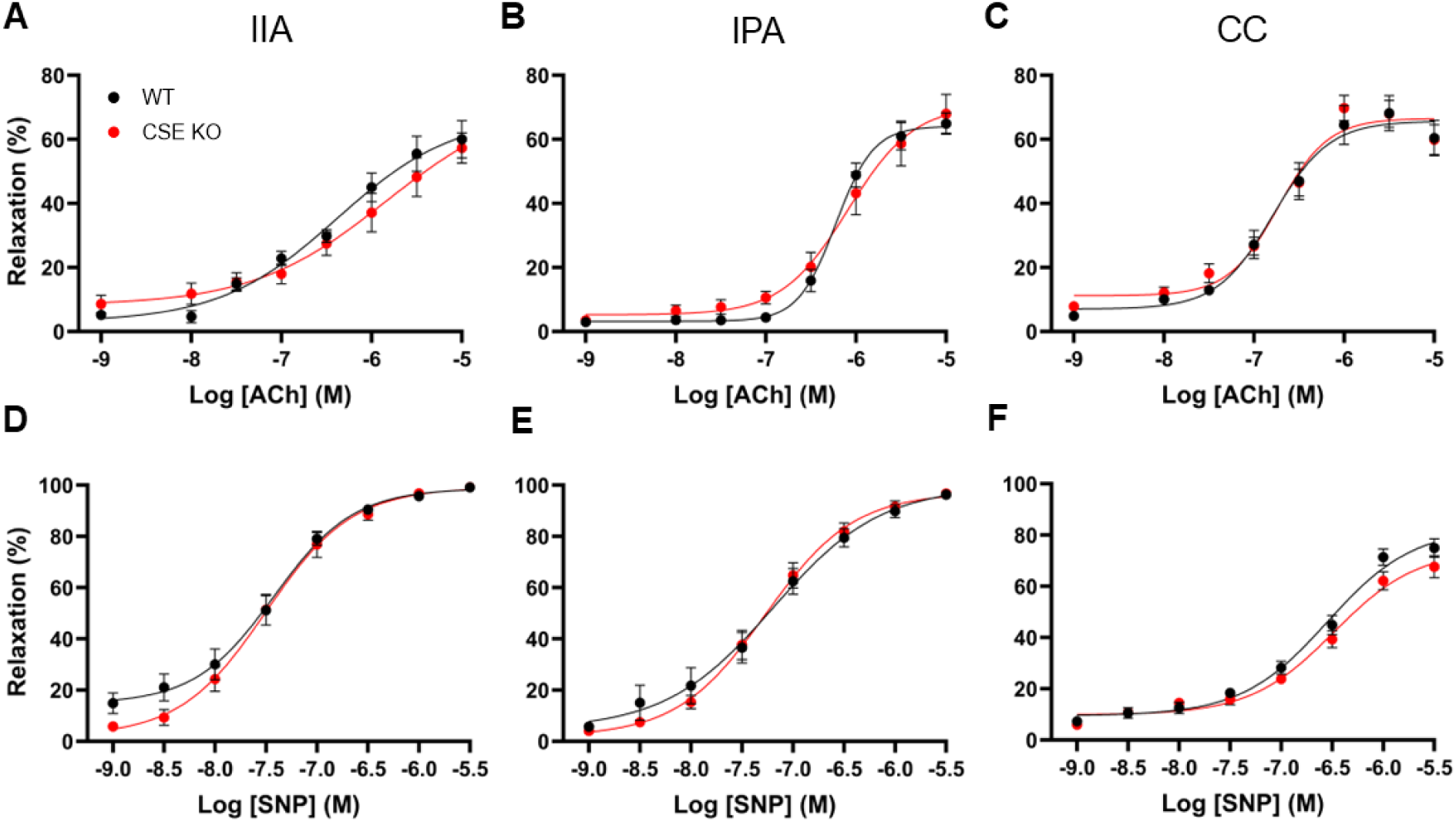
Evaluation of endothelium-dependent and -independent vasorelaxation from wild type (WT) and cystathionine γ-lyase knockout (CSE KO) mice. (A-C) Endothelium-dependent relaxation was assessed using cumulative doses of acetylcholine (ACh) in the internal iliac artery (IIA), internal pudendal artery (IPA), and corpus cavernosum (CC). (D-F) Endothelium-independent relaxation was evaluated using cumulative doses of sodium nitroprusside (SNP). Group differences were assessed by two-way repeated measures ANOVA, with no differences observed between groups. Data are expressed as mean ± SE for N = 13 mice per group.

**Figure 2.**
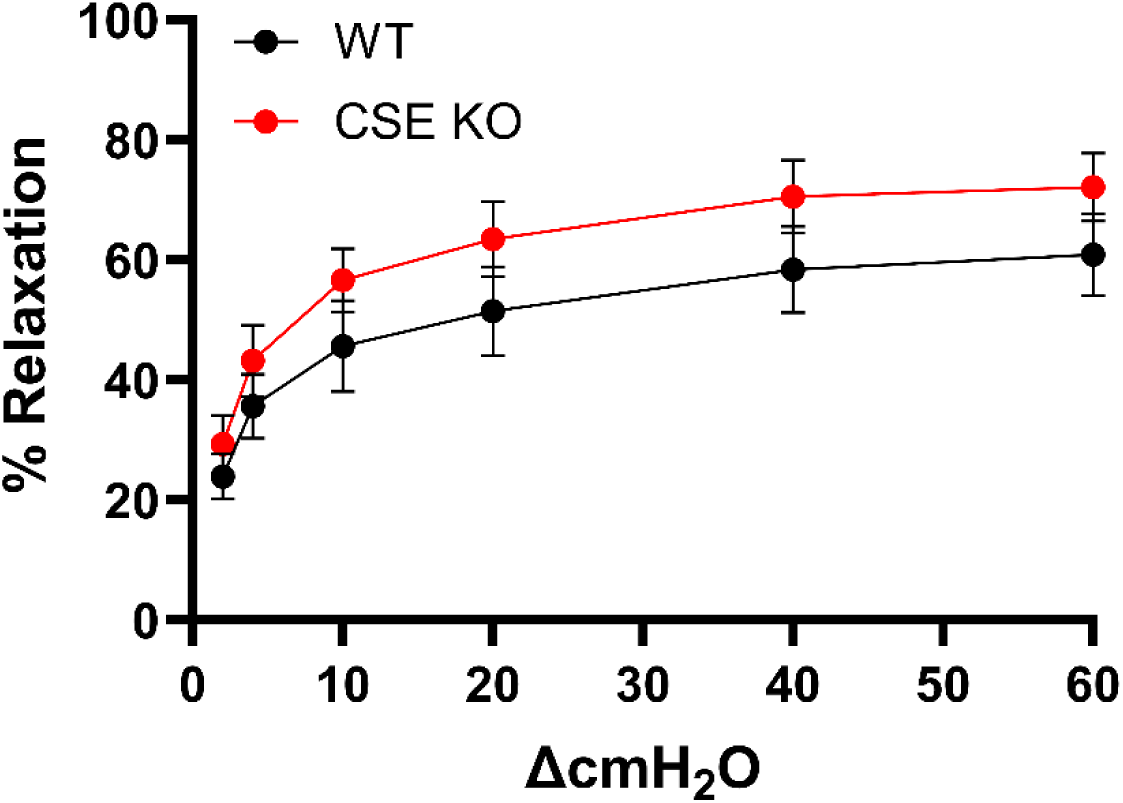
Endothelium-dependent flow-mediated vasodilation in the internal pudendal artery of wild type (WT) and cystathionine γ-lyase knockout (CSE KO) mice. Group differences were assessed by two-way repeated measures ANOVA, with no differences observed between groups. Data are expressed as mean ± SE for N = 9-10 mice per group.

### Neurogenic Control is altered in the CSE KO Corpus Cavernosum

To evaluate whether CSE deletion alters neurogenic-mediated vasodilation, electrical field stimulation (EFS) was applied in the presence of adrenergic and cholinergic blockers to assess non-adrenergic, non-cholinergic (NANC) relaxation. As demonstrated in Figure 3 (Panels A-C), the NANC-mediated relaxation in the IIA (p = 0.13) and IPA (p = 0.43) was comparable between WT and CSE KO mice. However, NANC-mediated relaxation in the CC was significantly reduced in CSE KO mice (p = 0.012), while Sidak’s post hoc testing revealed differences at stimulation frequencies of 4–16 Hz (p = 0.012, 0.004, 0.010, respectively).

**Figure 3.**
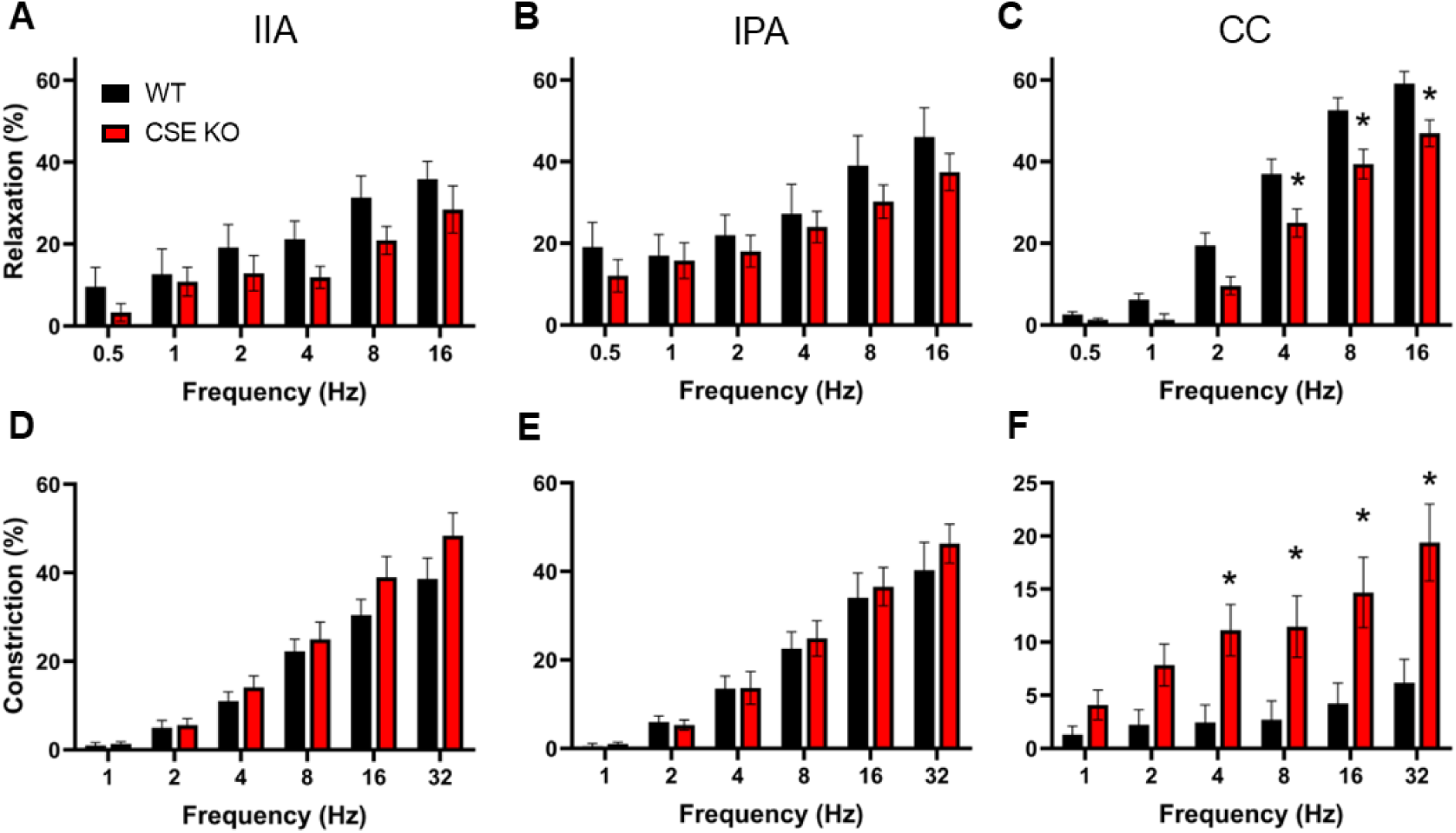
Evaluation of neurogenic vasorelaxation and vasoconstriction in erectile tissues from wild type (WT) and cystathionine γ-lyase knockout (CSE KO) mice. (A-C) non-adrenergic, non-cholinergic (NANC)-mediated relaxation was assessed in the internal iliac artery (IIA), internal pudendal artery (IPA), and corpus cavernosum (CC) using electrical field stimulation (EFS) following treatment with atropine and guanethidine. (D-F) Neurogenic contraction was evaluated in the IIA, IPA, and CC using increasing EFS frequencies. Group differences were assessed by two-way repeated measures ANOVA. *P < 0.05 at the indicated frequency for CSE KO vs. WT following Sidak’s multiple comparisons post-hoc test. Data are presented as mean ± SE for N = 13 mice per group.

To assess whether CSE deletion influences neurogenic contraction, EFS-induced vasoconstriction was measured in the absence of atropine, guanethidine, or pre-constriction. Figure 3 (Panels D-E) shows that neurogenic contraction was similar between WT and CSE-KO mice in the IIA (p = 0.27) and IPA (p = 0.66). However, EFS-induced contractility was significantly greater in the CC of CSE KO mice (p = 0.007), with Sidak’s post hoc testing indicating significantly heightened contractile responses to neurogenic stimulation at frequencies 4-32 Hz (p = 0.044, 0.041, 0.008, <0.001, respectively).

### α1-Adrenergic and Endothelin-1 (ET-1) Induced Vasoconstriction is Not Affected by CSE Deletion

To determine whether CSE deletion influences contractile responses to vasoconstrictors, cumulative dose-response curves for phenylephrine (PE, an α1-adrenergic agonist) and endothelin-1 (ET-1) were assessed. Figure 4A-C show PE-induced vasoconstriction in the IIA, IPA, and CC, with no significant differences observed between WT and CSE-KO mice (p > 0.05). Similarly, ET-1-induced contraction was also comparable between groups (p > 0.05, Figure 4D-F).

**Figure 4.**
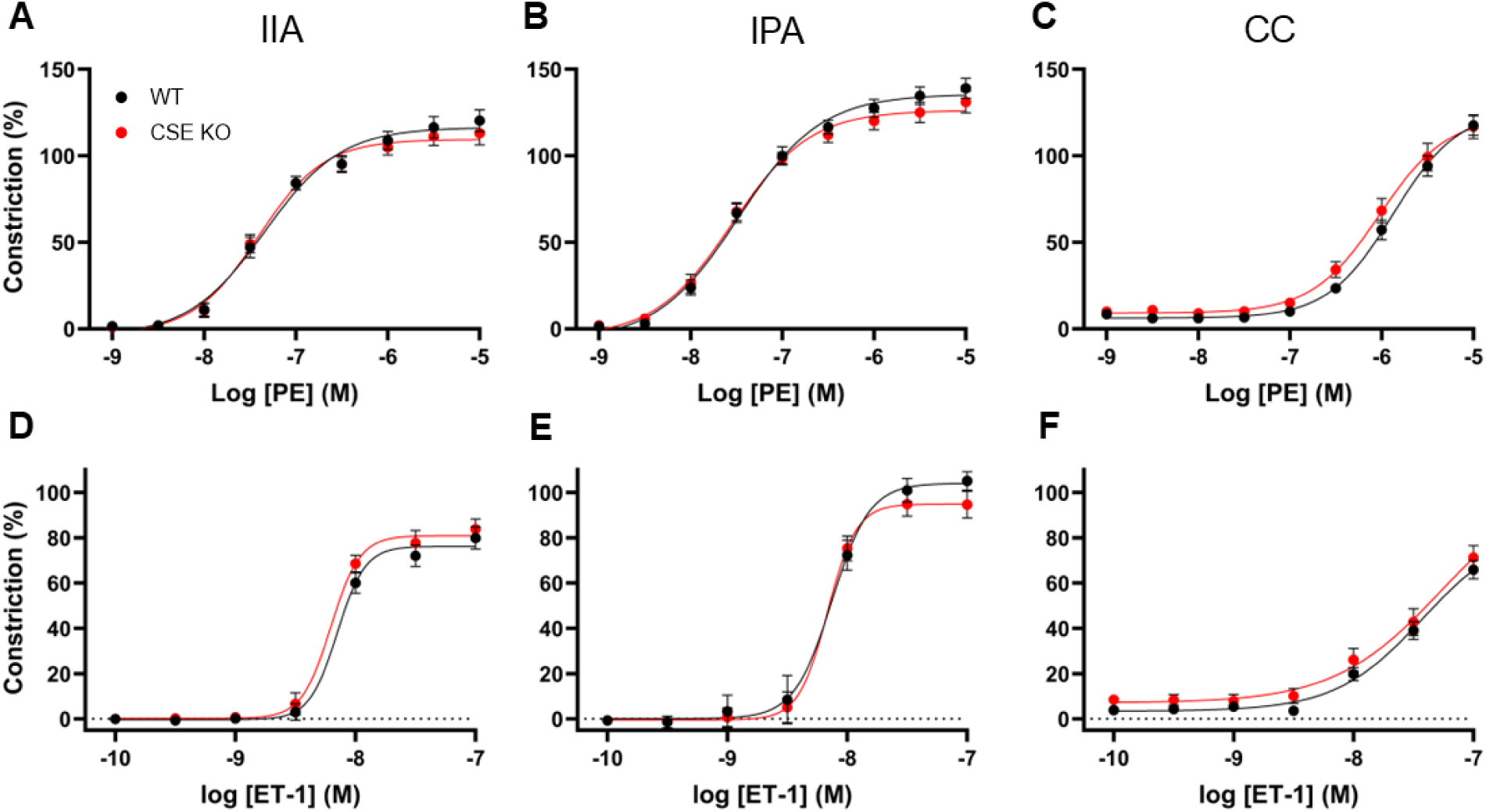
Evaluation of α1-adrenergic and endothelin-1 (ET-1)-induced vasoconstriction in erectile tissues from wild type (WT) and cystathionine γ-lyase knockout (CSE KO) mice. (A-C) α1-adrenergic vasoconstriction was assessed using cumulative doses of phenylephrine (PE) in the internal iliac artery (IIA), internal pudendal artery (IPA), and corpus cavernosum (CC). (D-F) Endothelin-1 (ET-1)-mediated vasoconstriction was assessed by cumulative dose application of ET-1. Group differences were assessed by two-way repeated measures ANOVA, with no differences observed between groups. Data are expressed as mean ± SE for N = 13 mice per group.

### TXA2 -Mediated Vasoconstriction is Increased in the Corpus Cavernosum

To assess whether CSE deletion alters TXA2 receptor-mediated vasoconstriction, cumulative dose-response curves to U-46619 were generated. As seen in Figure 5A-B, U-46619-induced contraction was not significantly different in the IIA (p = 0.25) and IPA (p = 0.61). However, U-46619-induced contraction was significantly greater in the CC of CSE-KO mice compared to WT (p = 0.016, Figure 5C).

**Figure 5.**
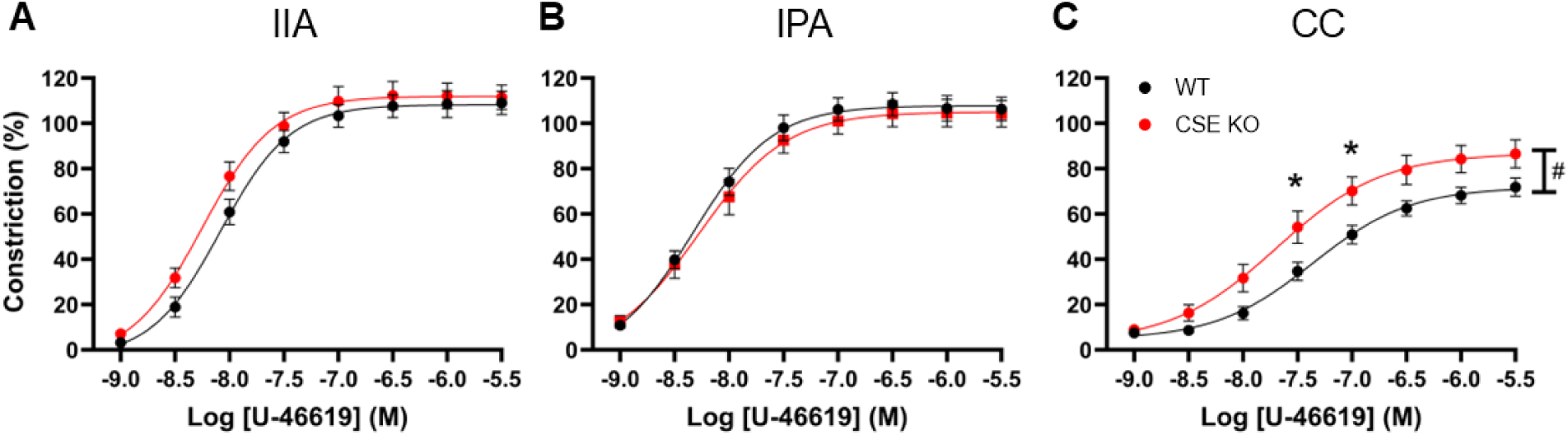
Evaluation of thromboxane A2 receptor-mediated vasoconstriction in erectile tissues from wild type (WT) and cystathionine γ-lyase knockout (CSE KO) mice. Cumulative doses of the thromboxane A2 receptor agonist U-46619 were applied to the A) internal iliac artery (IIA), B) internal pudendal artery (IPA), and C) corpus cavernosum (CC). Group differences were assessed by two-way repeated measures ANOVA. # main effect (P = 0.016) was observed between the groups. *P < 0.05 at the indicated concentration for CSE KO vs. WT following Sidak’s multiple comparisons post-hoc test. Data are presented as mean ± SE for N = 13 mice per group.

Sidak’s post hoc testing revealed significant increases in contraction at two doses in the middle of the dose-response curve (p = 0.019, 0.021, respectively).

### Vasodilatory sensitivity to H2S in erectile tissues is not altered by CSE deletion

To determine whether CSE deletion affects vascular responsiveness to exogenous H₂S, sodium sulfide (Na₂S), a rapid-releasing H₂S donor was applied to the IIA, IPA, and CC. As demonstrated in Figure 6, Na₂S-induced relaxation in the IIA (Panel A), IPA (Panel B), and CC (Panel C) was comparable between WT and CSE KO mice (p > 0.05), suggesting that smooth muscle reactivity to H₂S-mediated relaxation remains intact despite genetic deletion of this prominent H2S producing enzyme.

**Figure 6.**
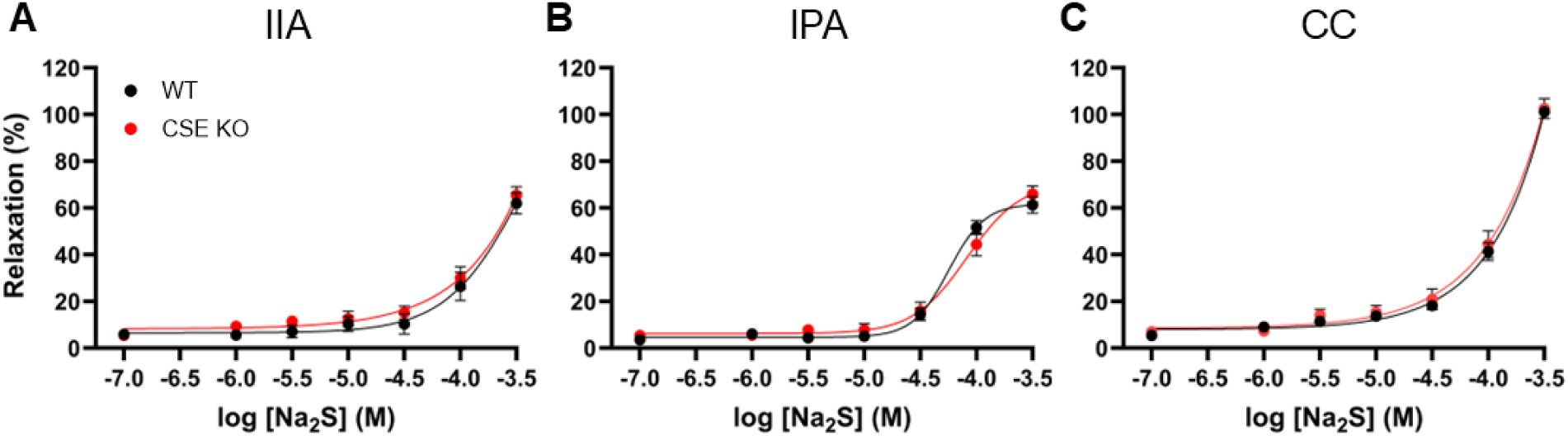
Evaluation of hydrogen sulfide (H2S)-mediated vasorelaxation in erectile tissues from wild type (WT) and cystathionine γ-lyase knockout (CSE KO) mice. Cumulative doses of the rapid-releasing H2S donor sodium sulfide (Na₂S) were applied to the A) internal iliac artery (IIA), B) internal pudendal artery (IPA), and C) corpus cavernosum (CC). Group differences were assessed by two-way repeated measures ANOVA, with no differences observed between groups. Data are expressed as mean ± SE for N = 13 mice per group.

### Histological Analysis of Vascular and Erectile Tissues

Masson’s Trichrome staining was performed to evaluate structural and fibrotic remodeling in the IIA, IPA, and CC tissue cross-sections. Representative images of IIA and IPA sections are presented in Figure 7. Quantification of lumen area, media area, and media-to-lumen area ratio for these arteries are reported in Table 1. There was no significant structural remodeling observed in these arteries with CSE deletion. Representative images of penile tissue cross-sections are presented in Figure 8. Fibrotic remodeling was assessed by the collagen-to-smooth muscle area ratio, whereby quantified data from the IIA, IPA, and CC are presented in Figure 9. We observed mean increases in the CSE KO condition of 27% and 21% in the collagen-to-smooth muscle area ratio for the IIA and IPA, respectively. However, these trends did not reach statistical significance (IIA: p = 0.093; IPA: p = 0.124). There were no differences between groups for collagen/smooth muscle in the CC (p = 0.471).

**Figure 7.**
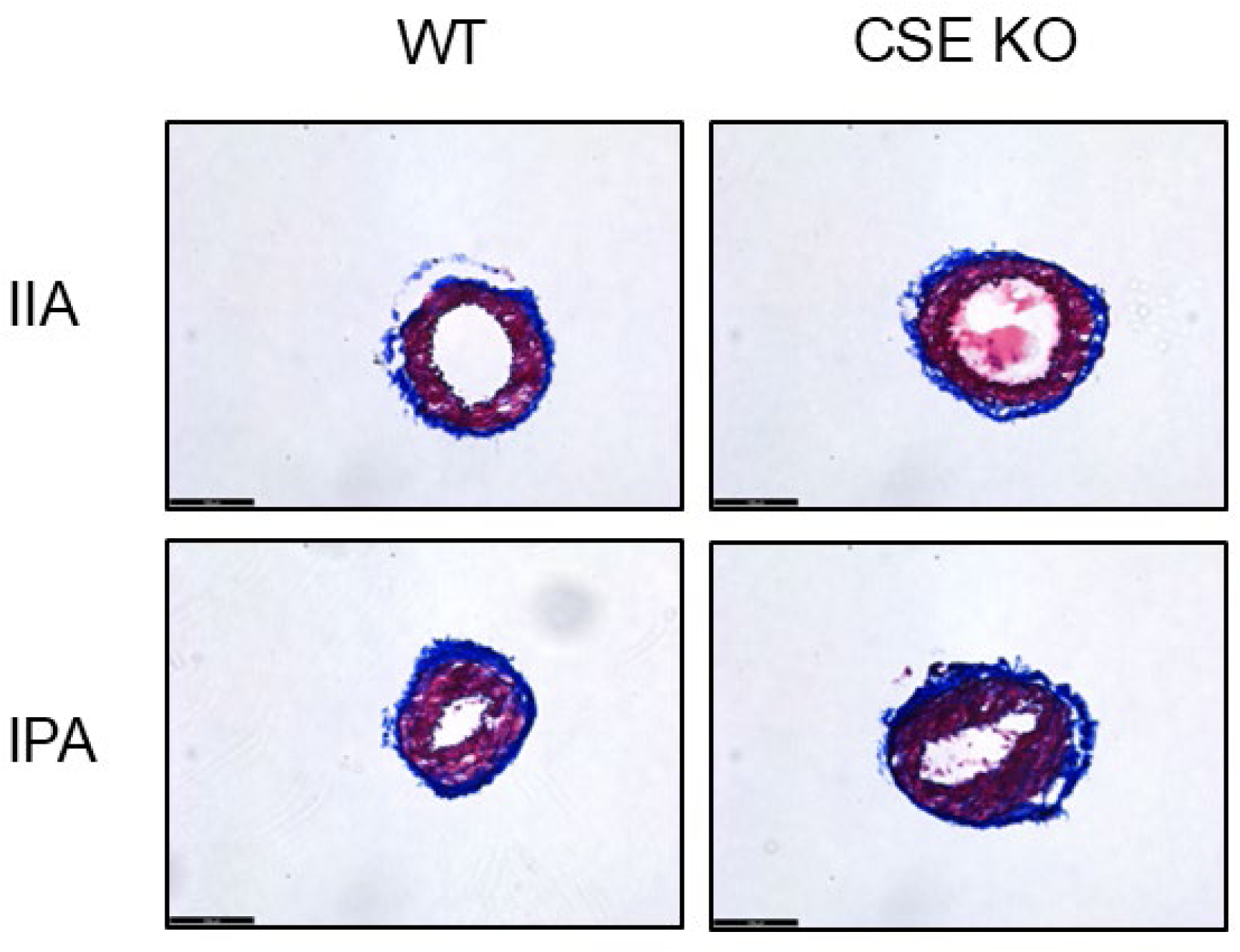
Representative histological sections of the internal iliac artery (IIA) and internal pudendal artery (IPA). Masson’s Trichrome staining was performed to assess vascular remodeling and presence of fibrotic remodeling in the IIA and IPA of wild-type (WT) and cystathionine γ-lyase knockout (CSE-KO) mice. Of note, arteries exhibit normal vascular architecture, with smooth muscle stained in red and collagen stained in blue. Images were taken at 20x magnification where the scale bar represents 100 μm.

**Figure 8.**
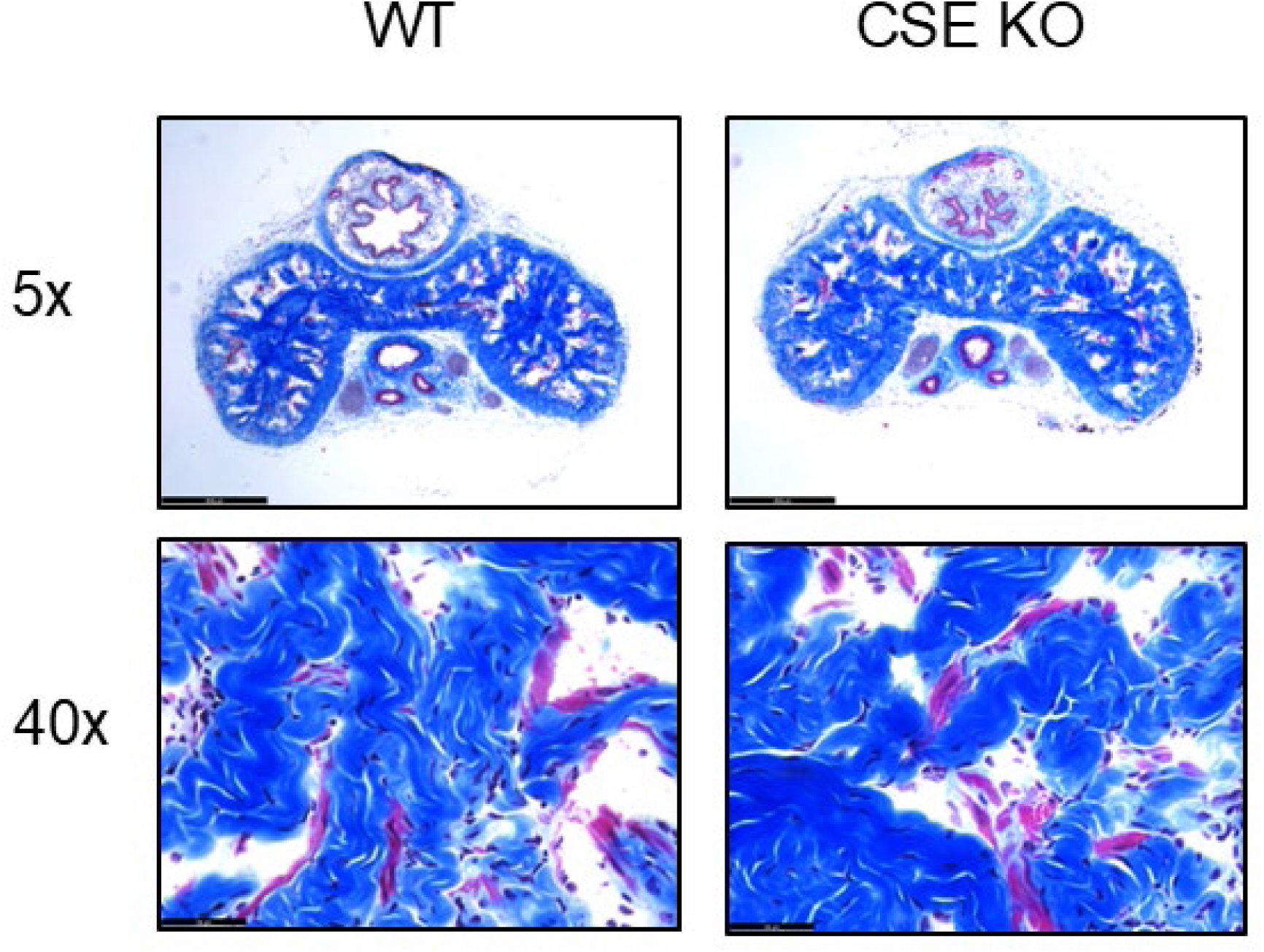
Histological assessment of penile tissue cross-sections. Masson’s Trichrome staining was used to evaluate collagen (blue) and smooth muscle (red) distribution in the corpus cavernosum (CC) of wild-type (WT) and cystathionine γ-lyase knockout (CSE KO) mice. Images of the whole tissue were taken at 5x magnification (top row) where the scale bar represents 500 µm. Zoomed in images of the CC (bottom row) were taken at 40x magnification where the scale bar represents 50 µm.

**Figure 9.**
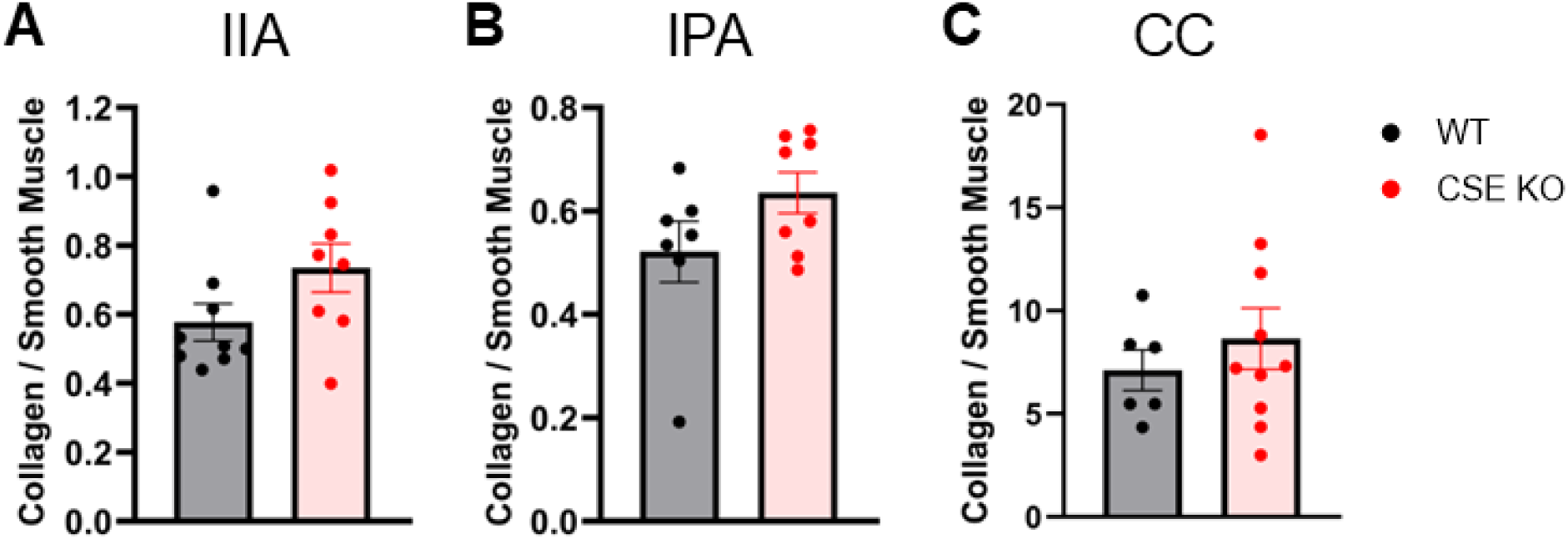
Quantitative histological analysis of fibrotic remodeling in erectile arteries and tissues. Smooth muscle and collagen contents were quantified from Masson’s trichrome stained images of wild type (WT) and cystathionine γ-lyase knockout (CSE KO) mice. Data are expressed as the collagen to smooth muscle ratio for the A) internal iliac artery (IIA), B) internal pudendal artery (IPA), and C) corpus cavernosum (CC). Group differences were assessed by student’s t-test. Data are presented as mean ± SE for N = 6-10 animals per group.

**Table 1.**
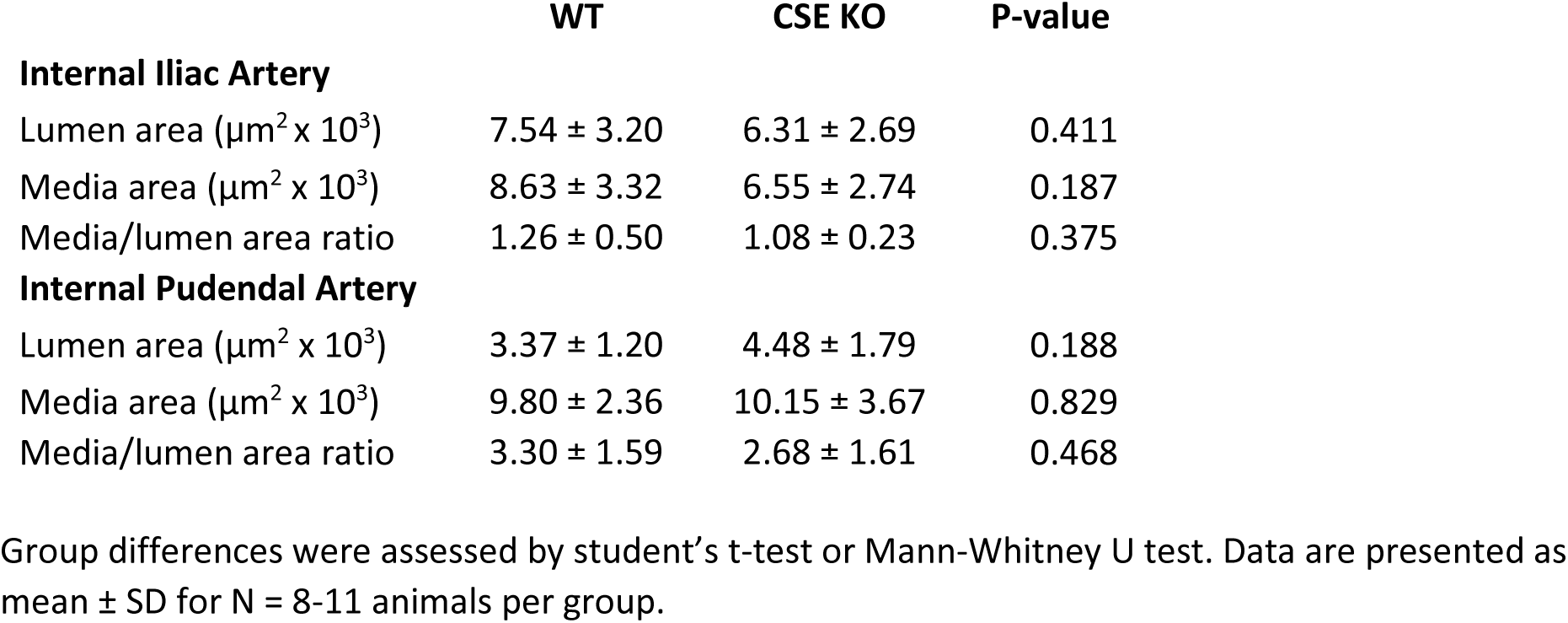
Vascular structure characteristics.

## DISCUSSION

The present study aimed to investigate the impact of chronic cystathionine γ-lyase (CSE) depletion on modulating neurovascular function pertinent to erectile physiology. By utilizing a CSE knockout (CSE KO) model, we assessed endothelial-dependent and -independent relaxation, NANC-nerve mediated vasodilation, and neurogenic vascular contractile responses in pre-penile arteries and erectile tissues. Our findings reveal that while CSE deletion does not impair nitric oxide (NO)-mediated relaxation or exogenous H₂S-induced vasodilation, it significantly alters neurogenic relaxation and contractility in the corpus cavernosum (CC) while enhancing thromboxane A2 receptor-mediated vasoconstriction. These results underscore the critical role of endogenous H₂S in maintaining erectile function and vascular homeostasis.

Our findings demonstrate that endothelium-dependent and endothelium-independent relaxation remain intact despite the absence of CSE (Fig. 1), as indicated by the preserved responses to acetylcholine (ACh) and sodium nitroprusside (SNP). Previous studies have shown that H₂S plays a vasodilatory role in multiple vascular beds, often acting synergistically with NO to modulate vascular tone (12, 34, 35). However, the lack of impairment in NO-mediated vasodilation in CSE KO mice suggests that compensatory mechanisms may sustain NO bioavailability in the absence of CSE-mediated H₂S. While H2S is typically reported to augment NOS efficiency and NO bioavailability, these two gasotransmitters do have a complicated relationship, and it is plausible that other endothelium derived hyperpolarizing factors upregulate in these erectile tissues when H2S availability is compromised (36).

Flow-mediated dilation (FMD) was also preserved in the IPA of CSE KO mice (Fig. 2), further indicating that the endothelial response to shear stress is maintained despite the loss of CSE-derived H2S. Our previous work shows that CSE KO mice exhibit elevated penile (24). H₂O₂ levels H2O2 has been identified to have vasodilatory actions in response to both ACh and shear stress in multiple resistance arteries (37–39). Gomez del Val et al. have recently demonstrated that mitochondria derived H2O2 from endothelial cells contribute to endothelium-dependent relaxations of penile arteries of high-fat diet fed rats, likely as a compensatory response to depressed NO production (40). While CSE is a cytosolic enzyme, it has been shown to produce H2S that can provide antioxidant effects in the mitochondria (41). It is therefore plausible that in the absence of CSE derived H2S, endothelial NO production may be depressed but compensated for by H2O2 or another hyperpolarizing factor.

Despite preserved endothelium-dependent relaxation, neurogenic-mediated erectile responses were significantly altered in the CC of CSE KO mice. Electrical field stimulation (EFS) revealed a reduction in non-adrenergic, non-cholinergic (NANC) relaxation, alongside heightened neurogenic contractility in the CC (Fig. 3). These findings suggest that H₂S plays a crucial role in modulating neurovascular signaling during penile erection. H2S has indeed been shown to exert a wide range of neuromodulatory actions (42). The erectile response is initiated by NANC-neurotransmission, of which nNOS derived NO is a predominant neurotransmitter (43). Given that the smooth muscle response to the NO donor SNP remained intact with CSE KO, the resulting impairment in NANC-mediated relaxation of the CC with CSE KO is therefore unlikely to be due to the smooth muscle response to NO and more likely due to altered neurotransmission. For instance, H2S has been shown to enhance NANC nerve outflow in the central vasculature through activation of the transient receptor potential ankyrin 1 (TRPA1) channel (44). In contrast, H2S has been shown to suppress sympathetic nerve transmission (45–47). This is particularly important in the erectile system, as the penis is primarily maintained in the flaccid state by sympathetic nerve activity (7). Our data further reveal that direct activation of the α1-adrenoceptor with phenylephrine is not changed in either of the arteries or the CC with CSE KO (Fig. 4a-c), indicating that the enhanced CC contraction with CSE KO in response to electrical stimulation was more likely an enhanced sympathetic transmission response. Moreover, ET-1 release from endothelial cells is another factor that contributes to maintaining the penis in a flaccid state. While we cannot verify that ET-1 release was unaltered in these tissues by H2S deficiency, the contractile response elicited by ET-1 was unaltered with CSE KO (Fig. 4d-f), removing sensitivity of the ET-1 receptor as a possible contributing factor to ED in the CSE null state.

Thromboxane A2 (TXA2) is a potent vasoconstrictor with a very short half-life that acts locally in its site of production. TXA2 is most notably produced by aggregated platelets but may also be produced by macrophages under conditions of inflammation, and by endothelial cells through the metabolism of arachidonic acid through the actions of cyclooxygenases and thromboxane synthases (48, 49). Elevated ROS levels, vascular damage and endothelial cell injury, and exposure of platelets to collagen all promote platelet aggregation and subsequent TXA2 production. We observed increased contractile responses in the CSE deficient corpus cavernosum to the TXA2 receptor agonist U-46619 (Fig. 5), indicating that any TXA2 produced in the CC in the CSE deficient state will have a greater impact on opposing the erectile response in vivo. Moreover, it is plausible that the elevated penile ROS that we observed with the CSE KO could lead to more TXA2 produced locally in the intact animal. Not only could this enhance smooth muscle contraction, it could also have a greater effect on opposing the dilatory effect of NO (50).

We observed no differences invoked by CSE KO in the relaxation responses to the fast-releasing H2S donor Na2S across any of the tissues investigated (Fig. 6), suggesting no changes in H2S sensitivity despite CSE deletion. However, we did observe much greater relaxations in the CC invoked by the high doses of Na2S relative to the two pre-penile arteries. This effect is the opposite of the NO donor SNP, where we observed more potent relaxations throughout the concentration range in both the IIA and IPA relative to the CC. Given this finding, combined with the other observed effects whereby CSE deficiency predominantly affected the corpus cavernosum, this work collectively suggests that H2S has a greater physiological impact on the CC than the pre-penile arteries of the IPA and the IIA. Extrapolating these observations, it is tempting to speculate that therapeutic H2S donors are more likely to impact the CC than these pre-penile arteries, at least in the absence of an overt disease stimulus. For instance, H2S is known to have anti-atherosclerotic effects, while the IPA is a highly atherosclerosis prone artery (51).

Under atherosclerotic conditions, either the absence of H2S or therapeutic H2S may have a greater influence on the physiology of the IPA. Stimulation of CSE expression via chronic histone deacetylase 6 inhibition has recently been found to reverse hypercholesterolemia-associated ED, although the relative contributions of these arteries involved in the erectile response remains unexplored (52).

Histological analysis of the IIA and IPA did not reveal significant structural differences in lumen or media area with CSE deficiency (Table 1). A trend toward increased collagen deposition was observed in these arteries of CSE KO mice (Figs. 7, 9), although these trends were not associated with any functional decline amongst any of the functional parameters that were tested. However, given that these trends were observed at a middle-age stage under controlled conditions, we cannot rule out the possibility that these trends could become exacerbated with advanced age or additional metabolic stressors. Loss of cavernous smooth muscle and increased cavernous collagen deposition is a hallmark of severe ED under several pathological conditions (21, 25). While we observed impaired erectile function in CSE KO mice (24), this impairment likely translates to more of a modest ED state than a severe one.

Despite the elevation in penile ROS levels with CSE deficiency, we did not observe any indication of increased fibrosis of the corpus cavernosum (Figs. 8, 9). It is thus unlikely that fibrosis or structural remodeling contributed to the physiological alterations observed in the CC or the impairment in erectile function observed in these mice.

## SIGNIFICANCE AND LIMITATIONS

This study provides important evidence that CSE-derived H₂S contributes to erectile function by modulating neurogenic control of the corpus cavernosum through relaxation and contraction mechanisms. Moreover, CSE-derived H2S protects the corpus cavernosum from hypersensitive vasoconstriction resulting from thromboxane A2 receptor activation. Given the increasing recognition of H₂S as a crucial vasoprotective molecule, our findings suggest that targeting the H₂S signaling pathway may offer therapeutic benefits for ED, particularly in conditions associated with altered neurotransmission. However, several limitations must be acknowledged. Our study focused exclusively on male mice, given that ED is a male-specific condition. However, H₂S is known to modulate vascular function more broadly, and future studies should explore whether similar vascular effects occur in other regions of the circulatory system, particularly in female reproductive physiology where vascular tone is also critical. Furthermore, we performed assessments at a single time-point which represents a middle-aged stage. The effects of prolonged H₂S suppression may become more evident at later stages associated with advanced age. Furthermore, our findings suggest that H₂S deficiency contributes to neurovascular dysfunction, while the precise molecular pathways underlying these effects remain unclear. Finally, the lack of significant vascular remodeling in the IIA and IPA suggests that more prolonged H₂S deficiency or additional stressors may be necessary to induce structural changes. Future studies should explore the effects of chronic metabolic stress, oxidative damage, or inflammation in CSE-deficient models to better understand the long-term consequences of H₂S deficiency on neurovascular and erectile health.

## CONCLUSION

This study highlights the critical role of CSE-derived H₂S in modulating penile vascular homeostasis. While endothelial-dependent and -independent relaxation remain preserved in the absence of CSE, neurogenic erectile responses are significantly impaired, and contractile sensitivity to TXA2 receptor activation is heightened, leading to increased vascular resistance in the corpus cavernosum. These findings provide mechanistic insight into the pathophysiology of erectile dysfunction associated with CSE deficiency and suggest that targeting the H₂S pathways may offer novel therapeutic opportunities for erectile disorders.

## DATA AVAILABILITY

Data will be made available upon reasonable request.

## AUTHOR CONTRIBUTIONS

Conceived and designed research : T.A.A, J.D.L; performed experiments: T.A.A, C.J.P., C.M.I.; analyzed data: T.A.A., C.J.P., C.M.I.; interpreted results of experiments: T.A.A., S.P.C., J.M.M-D., J.D.L.; prepared figures: T.A.A., C.M.I.; drafted manuscript: T.A.A.; edited and revised manuscript: S.P.C., J.M.M-D., J.D.L.; approved final version of manuscript: T.A.A., C.J.P., C.M.I., S.P.C., J.M.M-D., J.D.L.

